# Disequilibrium between *BRCA1* and *BRCA2* circular and messenger RNAs plays a role in breast cancer

**DOI:** 10.1101/2023.03.10.532029

**Authors:** C. Levacher, M. Viennot, A. Drouet, L. Beaussire, S. Coutant, JC Théry, S. Baert-Desurmont, M. Laé, P. Ruminy, C. Houdayer

## Abstract

Breast cancer is a frequent disease for which the discovery of markers for early detection or prognostic assessment remains challenging. Circular RNAs (circRNAs) are single-stranded structures in closed loops, produced by backsplicing. CircRNA and messenger RNA (mRNA) are generated cotranscriptionally and backsplicing and linear splicing compete against each other. As mRNAs are key players in tumorigenesis, we hypothesize that a balance disruption between circRNAs and mRNAs could promote breast cancer. Hence, we developed an assay for a simultaneous study of circRNAs and mRNAs, called Splice and Expression Analyses by exon Ligation and High Throughput Sequencing (SEALigTHS). Following SEALigHTS validation for *BRCA1* and *BRCA2*, our hypothesis was tested using an independent research set of 95 pairs of tumour and adjacent normal breast tissues. On this research set, ratios of *BRCA1* and *BRCA2* circRNAs/mRNAs were significantly lower in tumour breast tissue compared to normal tissue (p=1.6e-09 and p=4.4e-05 for *BRCA1* and *BRCA2*, respectively). Overall, we developed an innovative method to study linear and backsplicing, described the repertoire of *BRCA1* and *BRCA2* circRNAs, including 10 novel ones, and showed for the first time that a disequilibrium between *BRCA1* and *BRCA2* circRNAs and mRNAs plays a role in breast cancer.

## Introduction

Breast cancer (BC) represents ¼ of all cancers in women and ranks first in term of incidence among female cancers [1] (https://gco.iarc.fr). According to recent data from Global Cancer Statistics 2020, almost 700 000 women deaths (15.5% of all female deaths by cancer) are caused by BC [1]. BC is a heterogeneous disease with specific molecular, histological, and clinical features. Despite effective treatments, the need for new genetic markers for early detection or prognostic assessment remains. Recently, circular RNAs (circRNAs) have emerged as a promising field of study. CircRNAs are single-stranded covalent structures in closed loops without a 3’ cap or a 5’ poly-A tail, produced by backsplicing i.e. the junction of a donor site to an acceptor site located upstream [2] (supplementary figure 1). CircRNAs contain complete exons of protein coding genes, can be transcribed by the RNA polymerase II and mediated by the spliceosome. This circularization protects them from degradation by RNA exonucleases, their half-life is thus longer than their linear counterparts [3].

The expression of circRNAs is tissue and cell type specific [4,5] and several elusive roles have been described such as (i) microRNA (miRNA) sponges [6] or (ii) RNA Binding Protein sponges [7] or (iii) modulators of transcription [8]. In any case, previous work supports the implication of circRNAs in cell proliferation, apoptosis or metastasis [7,9,10]. With regards to BC, whole transcriptome-based studies showed deregulation of specific circular RNAs, pinpointing their role as miRNAs sponges within the frame of circRNA-miRNA-mRNA networks [10].

Given that circular RNA and mRNA are generated cotranscriptionally [11] and that the amount of circRNAs can exceed the amount of mRNA of the host gene [12], we believed that circRNA may have another, more pronounced impact on tumorigenesis by direct deregulation of mRNA levels. The involvement of mRNAs in cancer is indeed well documented e.g alternative splicing alterations are known to affect epithelial-mesenchymal transition, apoptosis and cell proliferation [13–16]. Moreover, pathogenic variations impacting splicing could represent the most frequent class of alterations in cancer [17]. Thus, any other class of transcripts that could impact mRNAs could influence the tumour phenotype and circRNA biogenesis has been precisely described to compete with pre-mRNA splicing in a muscleblind gene model [11]. We hypothesized that proper gene regulation could mandate a balance between both transcripts and balance disruption between circRNAs and mRNAs could promote tumorigenesis. To test this hypothesis, we used breast cancer samples and *BRCA1* and *BRCA2* genes as a model. To study messenger RNAs and circular RNAs without the complexity and high costs of transcriptomic RNA-seq approaches, we developed an alternative, innovative high-throughput and genes-targeted approach, called Splicing and Expression Analyses by exon Ligation and High Throughput Sequencing (SEALigHTS). SEALigHTS allowed the simultaneous study of splicing and backsplicing for *BRCA1* and *BRCA2* and measured the circRNAs/mRNAs balance.

We characterized novel *BRCA1* and *BRCA2* circular RNAs and described for the first time a disequilibrium in circRNA/mRNAs levels between tumour and normal breast tissues.

## Patients & Methods

### Patients

SEALigHTS was first validated on a formalin-fixed, paraffin-embedded (FFPE) set of samples from 72 selected female patients carrying germline pathogenic variations (PV) on *BRCA1* or *BRCA2* collected between 2017 and 2020. Written informed consent was obtained for all patients who were tested and diagnosed within the frame of genetic counselling. Hence, 164 FFPE tissue samples were obtained from these 72 patients, consisting of 23 pairs of tumours (invasive breast carcinomas consisting of all BC subtypes (12 Luminal, 2 HER2-positive and 9 triple-negative breast carcinomas (TNBCs)) and normal adjacent breast tissues, 32 normal mammary tissues from prophylactic mastectomy, 3 tumour serous invasive carcinoma and 41 normal ovarian tissues from prophylactic oophorectomy and 42 normal tissues from fallopian tubes (figure 1A). All samples were reviewed by an expert pathologist to confirm diagnosis and evaluate breast cancer subtype according to the following recommendations. Cases were designated ER-or PR-negative according to French national criteria if less than 10% of tumour cells expressed ER/PR [18]. Immunohistochemistry (IHC) was used to evaluate HER2 expression, with grading based on the American Society of Clinical Oncology (ASCO)/College of American Pathologists (CAP) criteria [19]. BC subtypes were defined as follows: tumours positive for either ER or PR and negative for HER2 were classified as luminal; tumours positive for HER2 were considered HER2-positive BC; tumours negative for ER, PR, and HER2 were considered as triple-negative BC (TNBC). Nine PV, identified in 15 patients should impact *BRCA1* or *BRCA2* splicing i.e. 4 single nucleotide variations (SNV), 3 large deletions, 2 large tandem duplications (Supplementary Table 1).

**Figure 1:**
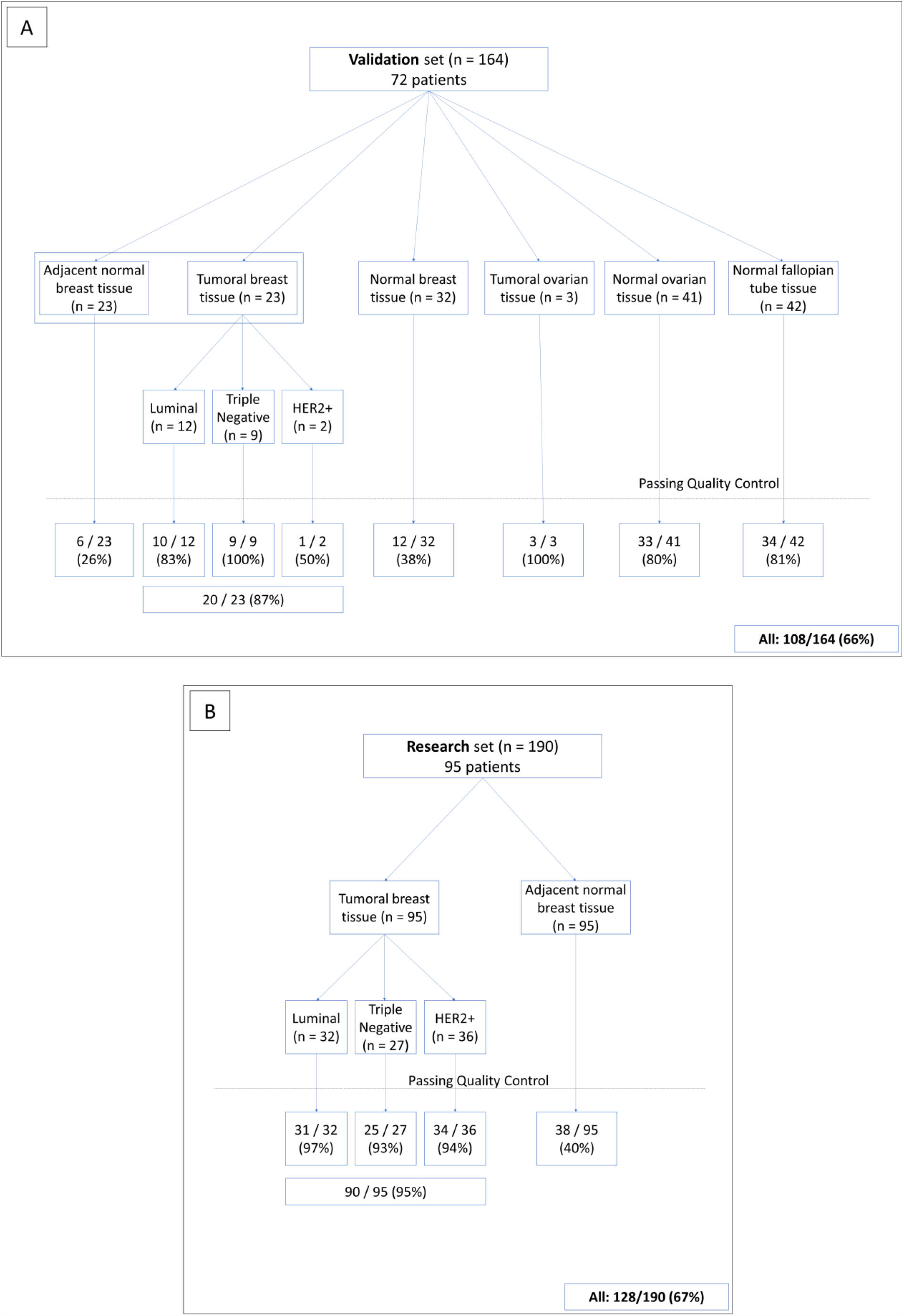
Description of the validation set (A) and the research set (B). The Quality Control corresponds to a number of UMI above 1500 for a given sample.

Following SEALigHTS validation, 190 normal and tumour breast tissues from 95 unselected, consecutively ascertained patients without genetic evaluation (*i*.*e*. they were not referred to the genetics clinics) were analysed in a research phase to test our hypothesis (figure 1B). Overall, 354 FFPE tissue samples were analysed. All FFPE tissues were retrieved from the biobank of the Henri Becquerel Cancer Center. Written informed consent was obtained for all patients.

### RNA extraction

One H&E-stained slide from formalin-fixed paraffin-embedded tissue (FFPE) was obtained for each sample and reviewed by an expert pathologist to evaluate tumour cellularity, which was always greater than 15%. RNA was then isolated from 8 consecutive 10-µm unstained slides using the automated Maxwell®16 Research extraction system (Promega, Madison, WI, USA) and the Maxwell®16 FFPE Plus LEV RNA Purification Kit following the manufacturer’s instructions, and were stored at -80°C. RNA concentration evaluation was performed by the Qubit fluorometer (Invitrogen, Carlsbad, CA).

### SEALigHTS assay

#### Principle

The SEALigHTS assay derives from RT-MLPseq [20] and LD-RTPCR [21], used to measure gene expression and to detect fusion transcripts [22–24] and was adapted to study simultaneously splicing and backsplicing. Briefly, probes designed at exon boundaries are joined when splicing occurs, then ligated and the resulting fragment is detected and quantified with a high throughput sequencer (figure 2A). All possible combinations of exons and their resulting transcripts, i.e. splicing and backsplicing, are amenable to detection and quantification. To search for allelic imbalance as an indirect consequence of nonsense mediated decay (NMD), probes targeting the 2 allelic versions of exonic single nucleotide polymorphisms (SNPs) were designed. Lastly, DNA contaminants were evaluated by using intronic probes.

**Figure 2:**
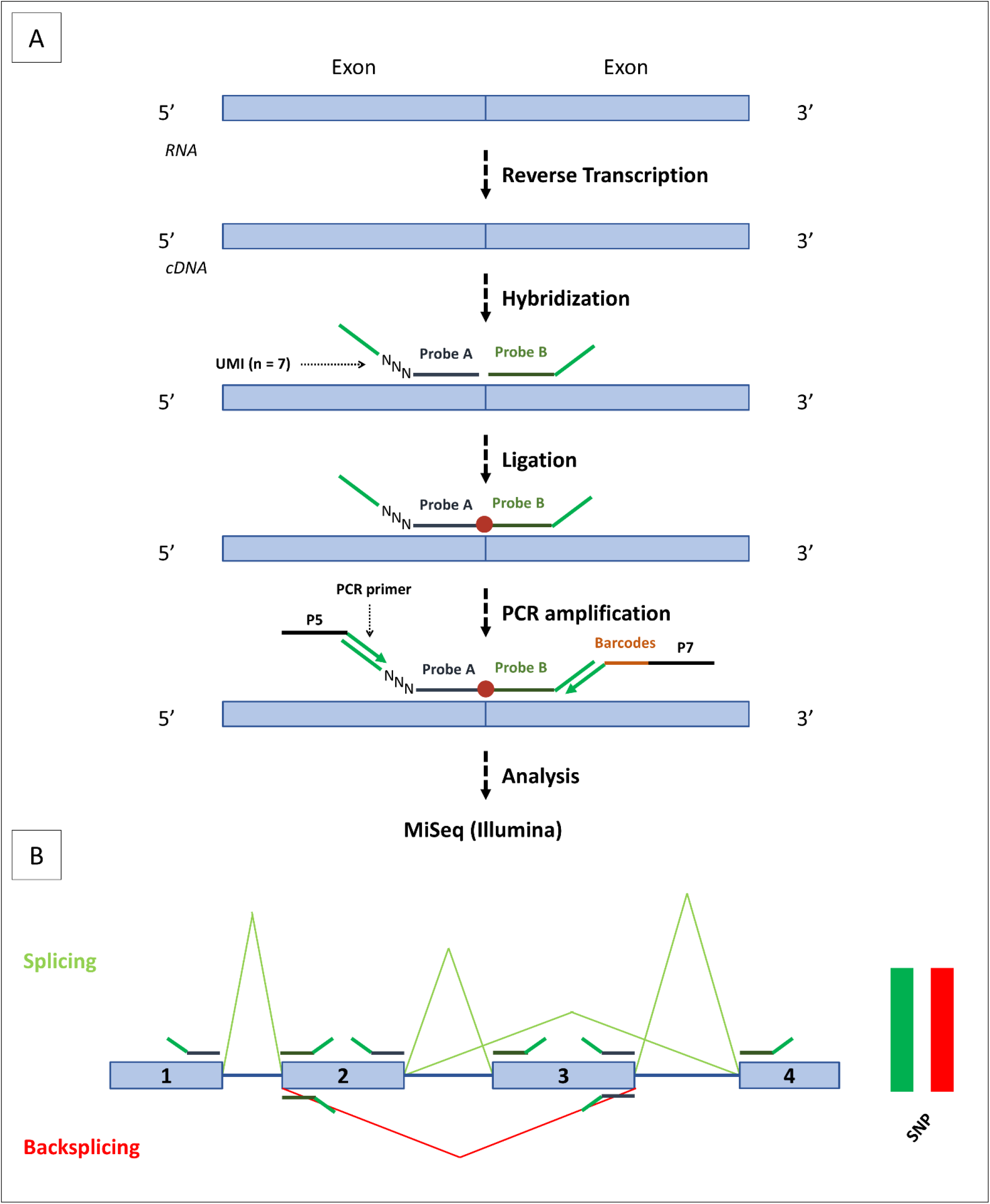
SEALIGHTS assay. (A) following cDNA synthesis, the probes are hybridized at exon extremities. Left probes include UMI consisting of a stretch of 7 random bases, depicted here as NNN. If hybridization takes place, probe ligation (depicted as a red dot) will occur. Then, PCR amplification will be performed using 2 universal complementary primers on the ends of each probe. Primers contain P5 or P7 adapters, which hybridise to the flowcell. Sample identification barcodes are included in the right primer. (B) Schematic representation of splicing and backsplicing results. Exons are represented as boxes and are numbered, with introns represented as lines. The probes are placed at exons extremities. Splicing junctions are represented in green, and backsplicing junctions in red. Peak heights depends on the number of UMI counts. Exon 3 skipping and exon3-exon 2 backsplicing are shown as examples. On the right side, a balanced heterozygous SNP (i.e. with the same number of UMI counts) is depicted in red and green for the 2 allelic versions.

#### Protocol

Oligonucleotides probes contained (i) specific sequences for each exon (20 – 30 bases length to obtain an optimal melting temperature of 70°C), (ii) Unique Molecular Identifiers (UMI), consisting of 7 random bases, to count the number of ligations (figure 2A). To study splicing and backsplicing, 114 probes were designed at exon boundaries for all *BRCA1* and *BRCA2* isoforms described in Davy and co-workers [25] and in the RJunBase, a web-accessible database of three types of RNA splice junctions (linear, back-splice, and fusion junctions) derived from RNA-seq data of non-cancerous and cancerous tissues [26]. To evidence NMD, we designed 30 probes assaying 6 and 4 SNPs on *BRCA1* and *BRCA2*, respectively. To control for DNA contamination, 3 intronic *BRCA1* probes were added in the probe mix (data available on request).

Following quantification, 0.2-600 ng of total RNA were converted into cDNA using *SuperScript™ VILO™ cDNA Synthesis* kit (Invitrogen, Carlsbad, CA). cDNAs were incubated 1h to 60°C with our mix of 147 oligonucleotides probes in 1X SALSA MLPA buffer (MRC Holland, Amsterdam, the Netherlands), ligated using the thermostable SALSA DNA ligase (MRC Holland, Amsterdam, the Netherlands) and amplified using barcoded primers containing P5 and P7 adaptor sequences with the *Q5 High-Fidelity 2X Master Mix* (NEB, Ipswich, MA). Amplification products were purified using AMPure XP beads (Beckman Coulter, Brea, CA) and analysed using a MiSeq sequencer (Illumina, San Diego, CA).

The reads are thus composed of at least 150 bases including the UMI, the left and right probes and the barcodes introduced during PCR amplification. Sequencing reads are demultiplexed using the barcodes, aligned with the sequences of the probes and exon junctions are counted. Quantitation is made thanks to the 7 random bases of the UMI, allowing 16 384 different combinations of unique molecules. Schematic backsplicing and splicing profiles were generated with a homemade Python script available on request (Figure 2B).

### Ratio circRNAs/mRNAs

Linear messenger RNAs (splicing) were distinguished from circular RNAs (backsplicing) thanks to the order of the probes. If a probe located at the 3’boundary of an exon is ligated with a probe located at the 5’boundary of the following exon, the transcript is linear. On the other hand, if this 3’ probe is ligated to a 5’probe of a preceding exon, the transcript is circular (Figure 2B). The relative proportion of a single circRNA on a gene is obtained by dividing the number of UMI of this circular RNA by the sum of UMI for every circular RNAs.

For each circular RNA, the circRNA/mRNA ratio is calculated by dividing the number of UMI of its circular junction by the average number of UMI of a linear junction of the gene:

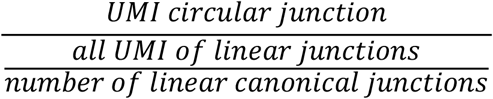

The overall circRNAs/mRNAs ratio is calculated by dividing the number of UMIs of all circular junctions by the number of UMI of all linear junctions of a gene:

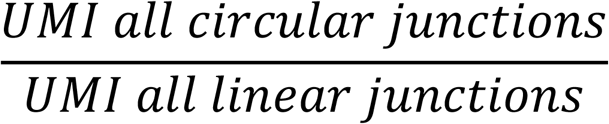

## Results

The first step was to validate the SEALigHTS assay for simultaneous detection of splicing and backsplicing, before embarking upon the research phase.

### General characteristics of SEALigHTS assay

Results were interpreted when at least 1500 different UMI per sample were counted for both *BRCA1* and *BRCA2*. The reason is that all known alternative transcripts above 1% were detected using this threshold. A total of 354 FFPE samples of various tissue types were analysed and 236 out of 354 samples (66.5%) passed the 1500 UMI threshold with an important disparity according to the type of sample (figure 1A). For breast tumour tissues in the validation set, 20 out of 23 (86%) samples met the quality criteria compared to 6 of the 23 (26%) samples from adjacent normal tissues. For prophylactic mastectomy tissues, 12 out of 32 (37.5%) samples passed the threshold whereas values were 33 of 41 (80.5%) for normal ovarian tissue samples and 34 of 42 (81%) for normal salpingian tissue samples. Overall, 108 samples were retained for the validation set. Similarly, for the breast tissues in the research set (figure 1B), 90 of the 95 (94.7%) tumour samples and 38 of the 95 (40%) normal tissue samples generated more than 1500 UMI, respectively. Thus, 128 samples were available for the test set.

### SEALigHTS assay validation

#### Splicing analyses

Sixty-six and 50 junctions making at total of 37 and 26 isoforms were correctly identified for *BRCA1* and *BRCA2*, respectively (Figures 3A and 4A). In our FFPE tissue material, SEALigHTS detected exon junctions above 0.26% and 0.15% for *BRCA1* and *BRCA2*, respectively as compared to RNAseq data from lymphoblastoid cell lines [25]. As a result, probe design was validated.

**Figure 3:**
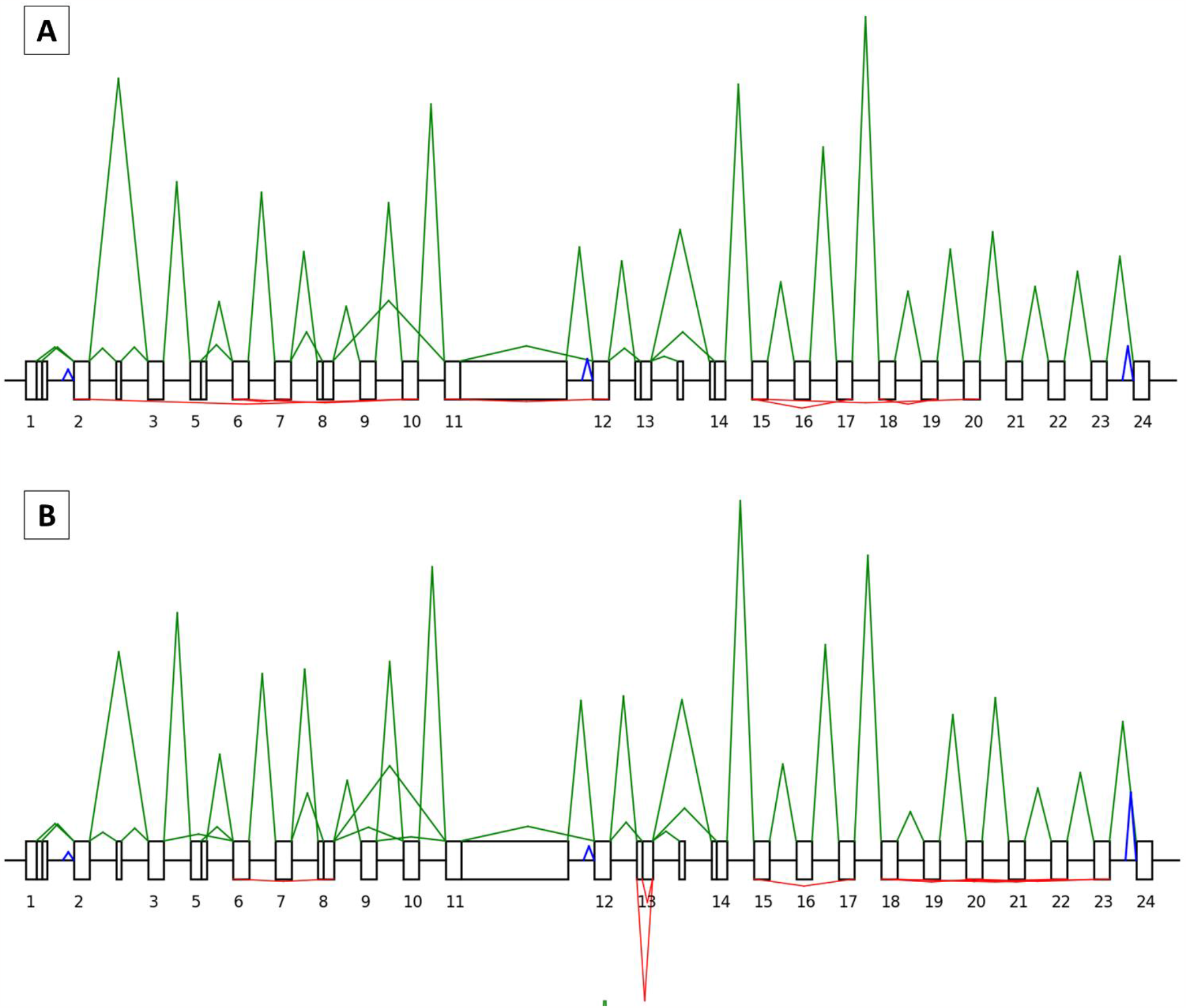
Representation of *BRCA1* splicing and backsplicing profiles. Exons are drawn as boxes and are numbered, with introns represented as lines. A solid line indicates alternative splice sites within exons. Non-numbered boxes correspond to alternative exons not included in the canonical transcript. Peak heights are relative to the number of UMI counts. For each profile, splicing is indicated in green above the boxes and backsplicing in red below the boxes. The blue peaks correspond to the signals of the intronic probes. **A** control profile (patient 20×0160, normal breast tissue): BRCA1 canonical transcript with different isoforms (delta 5q, delta 8p, delta 9-10, delta 11q, delta 13p, delta 14q) are shown. In red, below, backsplicing from exons 17 to 15 (circRNA_17-15), 19 to 18 (circRNA_19-18), 20 to 15 (circRNA_20-15), 7 to 6 (circRNA_7-6), 10 to 2 (circRNA_10-2) and 10 to 6 (circRNA_10-6). **B** tandem duplication of exon 13 (patient 20×0146, tumour breast tissue) detected thanks to two pseudo circular RNAs of exon 13 (both canonical and alternative 13p transcripts are produced), indicated with an arrow.

SEALigHTS identified the splicing consequences of genomic duplications, deletions, and SNV i.e. unexpected ligations between exons were evidenced. The *BRCA1* tandem duplication of exon 13 carried by patient 20×0146 was detected thanks to a pseudo circular RNA of exon 13 found in normal salpingian and ovarian tissue and in breast tumour tissue. This pseudo circular RNA results from the junction of the probe at the end of exon 13 to the probe at the beginning of the duplicated exon 13 (Figure 3B). Similarly, the *BRCA1* tandem duplication of exons 18 to 20 carried by patient 20×0131 resulted in a pseudo circular RNA joining exon 20 to duplicated exon 18 and was found in the patient’s breast cancer tissue sample.

Splicing analyses for patients 20×0133, 20×0180 and 20×0198, carrying a germline *BRCA1* deletion encompassing exons 8 to 13, led to an abnormal splice junction from exon 7 to exon 14, found in normal breast (Supplementary figure 2A), ovarian and salpingian tissues. In the breast tumour tissue, this abnormal skipping was found in combination with a decrease of the canonical junctions and an allelic imbalance, suggesting the loss of the wild type allele in the tumour (Supplementary figure 2B). Similarly, splicing analyses in patients 20×0129, 20×0184 and 20×0186, carrying a germline *BRCA1* deletion of exons 3 to 16, led to an abnormal splice junction from exons 2 to 17, evidenced in normal breast, ovarian and salpingian tissues. In the breast tumour tissue, the combination of abnormal skipping, decrease of the canonical junctions and allelic imbalance suggested the loss of the wild type allele. Lastly, the patient 20×0127 carrying a germline *BRCA1* deletion of exons 18 and 19 showed in both normal and tumour breast tissue an abnormal splice junction from exons 17 to 20.

Splicing anomalies caused by 3 distinct SNV were also detected. The *BRCA1* c.135-1G>C variation, leading to exon 5 skipping, was detected thanks to an abnormal junction of neighbouring exons in normal ovarian and salpingian tissues from patients 20×0172 and 20×0185. No allelic imbalance was observed at the heterozygous SNPs and the relative height of the peaks from the wild type junctions was similar to the height of the variant junction, in accordance with this in-frame exon skipping (Supplementary figure 3). The c.7805G>C variation in the *BRCA2* gene, leading to exon 16 skipping was detected with a junction between exon 15 and exon 17 in ovarian and salpingian tissue from patient 20×0175. The *BRCA2* c.67+3A>G variation (patient 20×0140) leading to exon 2 skipping was found thanks to an abnormal junction of exons 1 to 3, but an exon 1 to 4 junction was also evidenced in breast tumour tissue. This skipping of exons 2 and 3 has never been described before [25,26] and was not found in any other sample from our series.

In summary for the splicing outcome, SEALigHTS identified all expected splicing consequences of our selected variations, whether they were large deletions, duplications or point variations. More interesting, as analyses were done in the various tissues of interest, additional and tumour specific splicing anomalies were detected, as a plausible consequence of the second hit.

#### Backsplicing analyses

We counted 59 different physiological circular junctions for *BRCA1*. The complete description of circRNA landscape in breast tissues from the validation set is shown in Supplementary Table 2. We identified 10 novel ones not listed in RJunBase, i.e. *BRCA1*_circRNA_10-8; 12-6; 16-3; 18-2; 19-17; 20-20; 21-20; 23-22; 23-8p; 5q-3. For *BRCA2*, 23 circular junctions were detected, including 5 novel ones i.e. *BRCA2*_circRNA_13-12; 26-13; 6q-2; 7-5; 9-9 (Supplementary Table 3). On this selected set of samples, we found that *BRCA1* produced a higher amount of circular RNAs than *BRCA2*.

### Research set

#### Splicing study

Analysis of the research set of samples, consisting of pairs of normal and tumour breast tissues from unselected patients allowed us to found all the physiological mRNA junctions for the *BRCA1* and *BRCA2* genes previously identified in the validation set. In addition, in one patient, we found an abnormal junction from *BRCA2* exon 20 to exon 25 in the tumour breast sample, but not in the corresponding normal tissue. Relative peak height and absence of allelic imbalance were in accordance with an expected in frame skipping of exons 21 to 24 (Figure 4). No other splicing variations were found in the research set.

**Figure 4:**
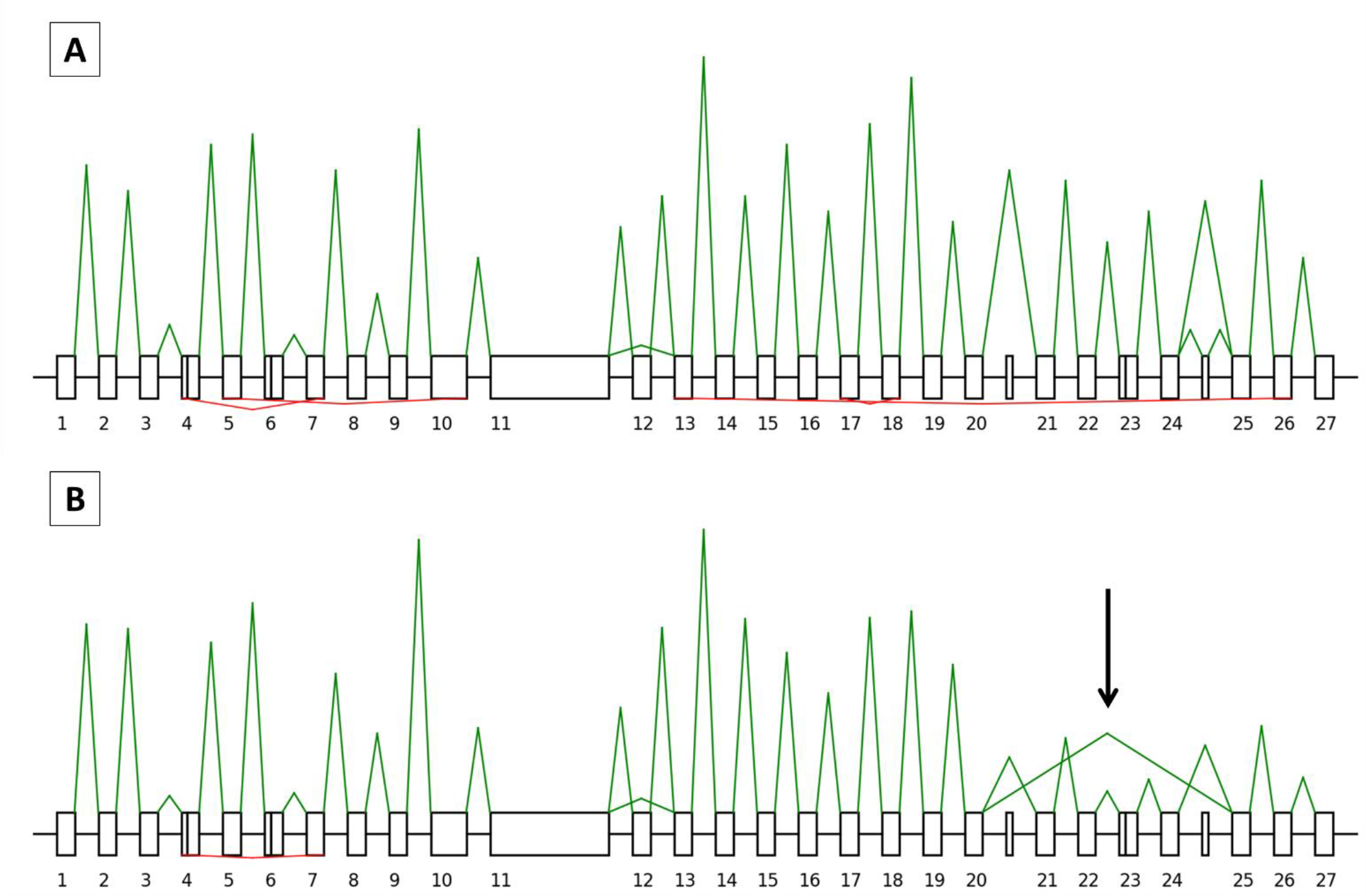
Representation of *BRCA2* splicing and backsplicing profiles for patient 20×0446, without a known pathogenic *BRCA2* germline variant. Exons are drawn as boxes and are numbered, with introns represented as lines. A solid line indicates alternative splice sites within exons. Non-numbered boxes correspond to alternative exons not included in the canonical transcript. Peak heights are relative to the number of UMI counts. For each profile, splicing is indicated in green above the boxes and backsplicing in red below the boxes. The blue peaks correspond to the signals of the intronic probes. **A** normal breast tissue, *BRCA2* canonical transcript with the delta 12 isoforms are shown. In red, below, backsplicing from exons 7 to 4 (circRNA_7-4), 10 to 5 (circRNA_10-5), 18 to 17 (circRNA_18-17) and 26 to 13 (circRNA_26-13). **B** tumour breast tissue, identification of an abnormal skipping from exon 20 to 25, indicated with an arrow, which was not present in the normal tissue.

#### Backsplicing study

All circular junctions identified in the validation set were found in the research set and we did not evidence other circRNAs. Considering both tumour and adjacent tissues, the ratio between circRNA and mRNA was 1.36 and 0.23 circular junctions per 100 linear junctions for *BRCA1* and *BRCA2*, respectively. In other words, *BRCA1* produced more circRNA than *BRCA2*. The 20 most frequent *BRCA1* and *BRCA2* circRNA found in normal and tumour tissues are indicated in Tables 1 and 2 and the full landscape of circRNAs detected is indicated in Supplementary Tables 4 and 5. Novel circRNAs are indicated Tables 3 and 4.

**Table 1:** The 20 most represented and novel *BRCA1* circular RNAs in the normal and tumour breast tissue of the research set. Twenty-one circRNA are listed as circRNA22-15 is among the 20 most represented in the tumour tissue, but not in the normal tissue. Alternatively, 5q-3 is among the 20 most represented in the normal tissue, but not in the tumour tissue. The novel circRNA are indicated in bold.

| Circular RNA name | *BRCA1* circular RNAs NORMAL |  |  | *BRCA1* circular RNAs TUMOUR |  |  |
| --- | --- | --- | --- | --- | --- | --- |
|  | Relative proportion to *BRCA1* circRNAs (%) | Frequency (%) (n= 38) | Ratio circRNA / mRNAs (%) | Relative proportion to *BRCA1* circRNAs (%) | Frequency (%) (n = 90) | Ratio circRNA / mRNAs (%) |
| BRCA1_circRNA_20-18 | 10,94 | 84,21 (32) | 4,73 | 11,18 | 92,22 (83) | 2,57 |
| BRCA1_circRNA_7-6 | 10,83 | 89,47 (34) | 4,68 | 7,93 | 87,78 (79) | 1,82 |
| BRCA1_circRNA_17-15 | 10,31 | 92,11 (35) | 4,46 | 9,49 | 92,22 (83) | 2,18 |
| BRCA1_circRNA_19-18 | 7,19 | 78,95 (30) | 3,11 | 5,80 | 84,44 (76) | 1,33 |
| BRCA1_circRNA_3-2 | 5,42 | 44,74 (17) | 2,34 | 2,17 | 46,67 (42) | 0,50 |
| BRCA1_circRNA_10-2 | 5,00 | 68,42 (26) | 2,16 | 3,78 | 72,22 (65) | 0,87 |
| BRCA1_circRNA_22-20 | 3,65 | 50 (19) | 1,58 | 4,81 | 87,78 (79) | 1,11 |
| BRCA1_circRNA_10-6 | 3,33 | 44,74 (17) | 1,44 | 2,66 | 63,33 (57) | 0,61 |
| BRCA1_circRNA_12-11 | 3,33 | 55,26 (21) | 1,44 | 5,25 | 84,44 (76) | 1,21 |
| BRCA1_circRNA_23-18 | 3,33 | 55,26 (21) | 1,44 | 5,15 | 81,11 (73) | 1,18 |
| BRCA1_circRNA_19-15 | 3,23 | 52,63 (20) | 1,40 | 2,59 | 72,22 (65) | 0,60 |
| BRCA1_circRNA_8-6 | 2,81 | 55,26 (21) | 1,22 | 2,08 | 56,67 (51) | 0,48 |
| BRCA1_circRNA_23-20 | 2,71 | 42,11 (16) | 1,17 | 5,51 | 81,11 (73) | 1,27 |
| BRCA1_circRNA_21-18 | 2,40 | 34,21 (13) | 1,04 | 2,68 | 63,33 (57) | 0,62 |
| BRCA1_circRNA_8-3 | 2,08 | 42,11 (16) | 0,90 | 2,91 | 76,67 (69) | 0,67 |
| **BRCA1_circRNA_5q-3** | 1,67 | 13,16 (5) | 0,72 | 0,71 | 22,22 (20) | 0,16 |
| BRCA1_circRNA_22-18 | 1,56 | 31,58 (12) | 0,68 | 2,37 | 61,11 (55) | 0,54 |
| BRCA1_circRNA_23-15 | 1,46 | 28,95 (11) | 0,63 | 1,24 | 43,33 (39) | 0,28 |
| BRCA1_circRNA_7-5 | 1,35 | 28,95 (11) | 0,59 | 1,69 | 41,11 (37) | 0,39 |
| BRCA1_circRNA_20-15 | 1,15 | 18,42 (7) | 0,50 | 1,75 | 55,56 (50) | 0,40 |
| BRCA1_circRNA_22-15 | 1,04 | 26,32 (10) | 0,45 | 1,69 | 57,78 (52) | 0,39 |

**Table 2:** The 20 most represented and novel *BRCA2* circular RNAs in the normal and tumour breast tissue of the research set. Twenty-one circRNA are listed as circRNA22-15 is among the 20 most represented in the tumour tissue, but not in the normal tissue. Alternatively, 5q-3 is among the 20 most represented in the normal tissue, but not in the tumour tissue. The novel circRNA are indicated in bold.

| Circular RNA name | *BRCA2 circular RNAs NORMAL* |  |  | *BRCA2 circularRNAs TUMOUR* |  |  |
| --- | --- | --- | --- | --- | --- | --- |
|  | Relative proportion to *BRCA2* circRNAs (%) | Frequency (%) (n= 38) | Ratio circRNA / mRNAs (%) | Relative proportion to *BRCA2* circRNAs (%) | Frequency (%) (n = 90) | Ratio circRNA / mRNAs (%) |
| BRCA2_circRNA_7-3 | 22,34 | 50 (19) | 2,76 | 26,13 | 76,67 (69) | 1,48 |
| BRCA2_circRNA_7-4 | 19,68 | 55,26 (21) | 2,43 | 21,18 | 76,67 (69) | 1,20 |
| BRCA2_circRNA_18-17 | 12,23 | 44,74 (17) | 1,51 | 12,61 | 61,11 (55) | 0,71 |
| **BRCA2_circRNA_7-5** | 6,91 | 21,05 (8) | 0,85 | 5,97 | 48,89 (44) | 0,34 |
| **BRCA2_circRNA_9-9** | 4,79 | 21,05 (8) | 0,59 | 2,10 | 24,44 (22) | 0,12 |
| **BRCA2_circRNA_26-13** | 4,26 | 21,05 (8) | 0,53 | 0,67 | 8,89 (8) | 0,04 |
| BRCA2_circRNA_21-20 | 3,72 | 7,89 (3) | 0,46 | 1,51 | 18,89 (17) | 0,09 |
| BRCA2_circRNA_13-11 | 3,19 | 13,16 (5) | 0,39 | 7,39 | 47,78 (43) | 0,42 |
| BRCA2_circRNA_4-3 | 3,19 | 13,16 (5) | 0,39 | 3,61 | 32,22 (29) | 0,20 |
| BRCA2_circRNA_14-13 | 2,66 | 13,16 (5) | 0,33 | 1,43 | 14,44 (13) | 0,08 |
| BRCA2_circRNA_10-4 | 2,13 | 7,89 (3) | 0,26 | 1,01 | 10 (9) | 0,06 |
| **BRCA2_circRNA_13-12** | 2,13 | 5,26 (2) | 0,26 | 0,76 | 10 (9) | 0,04 |
| **BRCA2_circRNA_6q-2** | 2,13 | 10,53 (4) | 0,26 | 0,76 | 10 (9) | 0,04 |
| BRCA2_circRNA_10-5 | 1,60 | 7,89 (3) | 0,20 | 0,67 | 7,78 (7) | 0,04 |
| BRCA2_circRNA_10-8 | 1,60 | 7,89 (3) | 0,20 | 1,43 | 17,78 (16) | 0,08 |
| BRCA2_circRNA_11-11 | 1,60 | 7,89 (3) | 0,20 | 1,18 | 13,33 (12) | 0,07 |
| BRCA2_circRNA_10-3 | 1,06 | 5,26 (2) | 0,13 | 1,01 | 12,22 (11) | 0,06 |
| BRCA2_circRNA_19-17 | 1,06 | 5,26 (2) | 0,13 | 3,28 | 32,22 (29) | 0,19 |
| BRCA2_circRNA_24-19 | 1,06 | 5,26 (2) | 0,13 | 2,10 | 20 (18) | 0,12 |
| BRCA2_circRNA_24-22 | 1,06 | 5,26 (2) | 0,13 | 1,76 | 16,67 (15) | 0,10 |
| BRCA2_circRNA_24-21 | 0,53 | 2,63 (1) | 0,07 | 2,44 | 28,89 (26) | 0,14 |

**Table 3:** Novel *BRCA1* circular RNA found RNAs in the normal and tumour breast tissue of the research set.

| Circular RNA name | *BRCA1 novel circular RNAs NORMAL* |  |  | *BRCA1 novel circular RNAs TUMOUR* |  |  |
| --- | --- | --- | --- | --- | --- | --- |
|  | Relative proportion to *BRCA1* circRNAs (%) | Frequency (%) (n= 38) | Ratio circRNA / mRNAs (%) | Relative proportion to *BRCA1* circRNAs (%) | Frequency (%) (n = 90) | Ratio circRNA / mRNAs (%) |
| **BRCA1_circRNA_5q-3** | 1,67 | 13,16 (5) | 0,72 | 0,71 | 22,22 (20) | 0,16 |
| **BRCA1_circRNA_23-8p** | 1,15 | 23,68 (9) | 0,50 | 0,76 | 40 (36) | 0,18 |
| **BRCA1_circRNA_19-17** | 1,04 | 21,05 (8) | 0,45 | 0,85 | 38,89 (35) | 0,19 |
| **BRCA1_circRNA_18-2** | 0,63 | 13,16 (5) | 0,27 | 0,25 | 14,44 (13) | 0,06 |
| **BRCA1_circRNA_16-3** | 0,52 | 13,16 (5) | 0,23 | 0,07 | 4,44 (4) | 0,02 |
| **BRCA1_circRNA_23-22** | 0,42 | 10,53 (4) | 0,18 | 0,10 | 6,67 (6) | 0,02 |
| **BRCA1_circRNA_10-8** | 0,31 | 7,89 (3) | 0,14 | 0,66 | 27,78 (25) | 0,15 |
| **BRCA1_circRNA_20-20** | 0,31 | 5,26 (2) | 0,14 | 0,41 | 11,11 (10) | 0,09 |
| **BRCA1_circRNA_12-6** | 0,10 | 2,63 (1) | 0,05 | 0,19 | 12,22 (11) | 0,04 |
| **BRCA1_circRNA_21-20** | 0,10 | 2,63 (1) | 0,05 | 0,10 | 6,67 (6) | 0,02 |

**Table 4:** Novel *BRCA2* circular RNA found RNAs in the normal and tumour breast tissue of the research set.

| Circular RNA name | *BRCA2 circular RNAs NORMAL* |  |  | *BRCA2 circular RNAs TUMOUR* |  |  |
| --- | --- | --- | --- | --- | --- | --- |
|  | Relative proportion to *BRCA2* circRNAs (%) | Frequency (%) (n= 38) | Ratio circRNA / mRNAs (%) | Relative proportion to *BRCA2* circRNAs (%) | Frequency (%) (n = 90) | Ratio circRNA / mRNAs (%) |
| **BRCA2_circRNA_7-5** | 6,91 | 21,05 (8) | 0,85 | 5,97 | 48,89 (44) | 0,34 |
| **BRCA2_circRNA_9-9** | 4,79 | 21,05 (8) | 0,59 | 2,10 | 24,44 (22) | 0,12 |
| **BRCA2_circRNA_26-13** | 4,26 | 21,05 (8) | 0,53 | 0,67 | 8,89 (8) | 0,04 |
| **BRCA2_circRNA_13-12** | 2,13 | 5,26 (2) | 0,26 | 0,76 | 10 (9) | 0,04 |
| **BRCA2_circRNA_6q-2** | 2,13 | 10,53 (4) | 0,26 | 0,76 | 10 (9) | 0,04 |

We identified 59 circRNAs for *BRCA1* and 23 for *BRCA2*. As shown in Tables 1 and 2 and supplementary Tables 4 and 5, their frequency varied whatever the tissue, with almost ubiquitous (found in 32/38 samples) to rare (found in one sample) circRNAs. The diversity of their repertoire ranged from 11,18 % to 0.07% for *BRCA1* and 26.13% to 0.34% for *BRCA2* and is less pronounced for *BRCA2* as circRNAs_7-3 and 7-4 accounted for half of the total *BRCA2* circRNAs. As compared to mRNAs, the maximum ratio is 4.73% for BRCA1_circRNA_20-18, in other words this circRNA made 4.7% of a mean mRNA junction for *BRCA1* and is found 4.73 times for 100 *BRCA1* mRNA molecules. The same circRNAs were identified in both normal and tumour breast tissues. However, individual variations occurred, the most pronounced were i) for *BRCA1* circRNA_23-20 with an increase in relative proportion from 2.71% in the normal tissue to 5.51% in the tumour tissue and circRNA_3-2 with a decrease from 5.42% in the normal tissue to 2.17% in the tumour tissue ii) for *BRCA2*, the novel circRNA 26-13 dropped from 4.26% in the normal tissue to 0.67% in tumour tissue. At the exception of *BRCA1* circRNA_23-20 (see above), lower circRNA/mRNA ratio were observed for all *BRCA1* and *BRCA2* circRNAs, which prompted us to calculate the overall circRNAs/mRNAs ratio.

For *BRCA1*, the average ratio circRNAs/mRNAs was significantly lower in tumour compared to normal adjacent mammary tissue (1.14 vs 1.89, p-value= 1.6e-09) (figure 5A). This held true for *BRCA2* (0.23 vs 0.51, p-value= 4.4e-05) (figure 5B). We then looked at the histological subtype and for both *BRCA1* and *BRCA2*, no significant difference in the circRNAs/mRNAs ratio was found for the 3 BC subtypes (luminal, TNBC and HER2+) (figure 5C and 5D). Similarly, no significant difference was found between grade II and grade III samples (*BRCA1*: 1.18 vs 1.11, p-value= 0.47; *BRCA2*: 0.24 vs 0.2, p-value=0.33) (figure 5E and 5F). Overall these results showed a circRNA/mRNA disequilibrium which was not explained by histological subtype and proliferation status.

**Figure 5:**
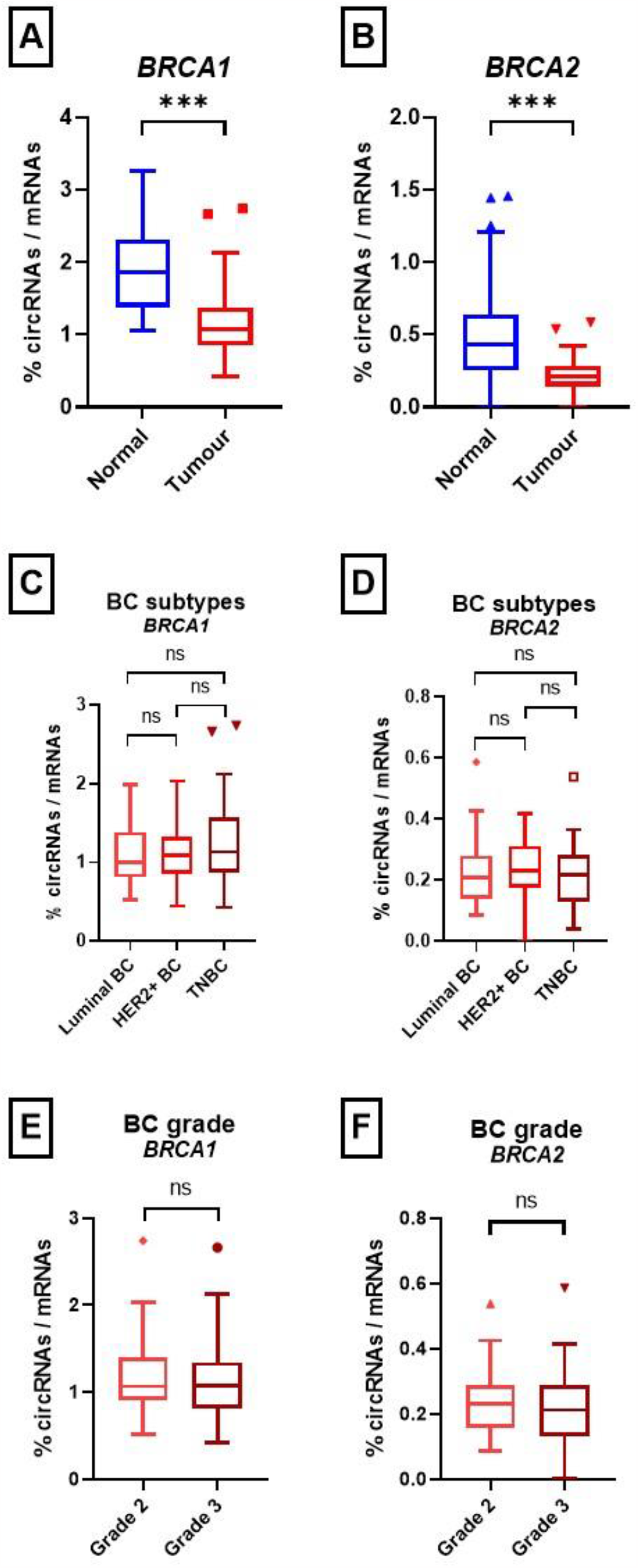
circular RNAs/messenger RNAs ratios. Ratios for normal and tumour tissues are indicated for *BRCA1* (A) and *BRCA2* (B). Ratios according to histological subtype (Luminal, HER2+ and Triple Negative Breast Cancer: TNBC) for *BRCA1* (C) and *BRCA2* (D). Ratios according to grade are indicated for *BRCA1* (E) and *BRCA2* (F). (ns: non-significant; ***: p < 0.001; Student test).

## Discussion

Firstly, our study provides the community with an innovative, simple and high throughput method for simultaneous detection of splicing and backsplicing, validated and applied in a series of 354 FFPE samples. Secondly, we demonstrated for *BRCA1* and *BRCA2* a disequilibrium in the circRNA/mRNA ratio between tumour and normal breast tissues. These two aspects will be discussed consecutively.

SEALigHTS simultaneously detects mRNAs and circRNA thanks to a simple design of probes located at exon boundaries and following ligation when splicing and backsplicing occur. This assay requires 2 days for the whole procedure i.e. reverse transcription, hybridization of the probes, ligation, PCR amplification and analysis on a high throughput sequencer. No specific material or platform are needed, SEALigHTS could be implemented in any laboratory with next-generation sequencing facilities. In our hands, the cost per sample was 30 euros (#30 USD). Another strong asset is the low quality and quantity of input RNA needed, as little as 70 ng, as demonstrated by the variety of tissue assayed (breast, ovarian and salpingian). With a quality threshold of 1500UMI, more than 90% of the tumour samples were successfully analysed. Low cellularity is obviously a limit and only 37.5% of normal breast tissue samples provided us with reliable data. It could be explained by the fact that breast tissue has a lower cellularity compared to breast tumour as it is mainly composed of adipocytes, fibrosis and rare duct and lobules. Nevertheless, our ability to handle FFPE material suggests that a wide variety of tissues could be robustly assayed. Moreover, the length of cDNA template is not an issue because probes need only about 60 bases long template to be hybridized. Joint analysis of messenger RNAs and circular RNAs is made easy since no treatment of the sample with exoribonucleases is required and the same bioinformatics tool is used for both transcripts. In theory, SEALigHTS can detect all kinds of splicing events, directly or indirectly. Hence, we successfully identified alternative splicing events and splicing consequences of SNV, large deletions and duplications from the validation set and even demonstrated additional events e.g. second hits in tumour RNA from both datasets. Although this is a targeted approach, intronic exonization should be detected as a decrease in the number of ligations is expected at the corresponding exon junctions. Allelic imbalance is detected by using exonic SNPs, and more largely we believe that gene expression could be calculated from the splicing data, as RT-MLPseq and LD-RT-PCR provided robust gene expression measurements [21,22].

SEALigHTS deciphered *BRCA1* and *BRCA2* backsplicing landscape in normal and tumour breast tissues and identified already described circRNA, but also 10 and 5 novel ones for *BRCA1* and *BRCA2*, respectively. Previous literature on breast cancer suggested that specific circular RNAs would be deregulated between normal and tumour tissues [27,28], but despite the fact that *BRCA1* and *BRCA2* are master genes of homologous recombination and breast cancer predisposition, our study is the first to address the question of their circular RNAs in breast cancer. We first evidenced a larger number of circRNAs in *BRCA1* than in *BRCA2*, in accordance with RJunBase, with a similar qualitative repertoire between normal and tumour tissues. We did show individual circRNA variations e.g. *BRCA1*_circRNA_3-2 and *BRCA2*_circRNA_26-13, that could be linked to the tumour process but these findings should be replicated on a larger series to draw definite conclusions. Hence, if a functional importance for *BRCA1*_circRNA_3-2 could be speculated because *BRCA1* exon 2 contains the wild type translation initiation sequence, a putative functional mechanism for *BRCA2* circRNA_26-13, which nearly disappears from tumour samples, remains unknown. Moreover, the relative proportion of each circRNA remains low as compared to its mRNA counterpart and do not exceed 5%.

On the other hand, we demonstrated, for the first time, a decrease of 40% and 55% in the circRNA/mRNA ratio for *BRCA1* and *BRCA2*, respectively, in breast cancer tissues compared to normal adjacent tissues. This disequilibrium can be viewed as a cause or a consequence of tumorigenesis. circRNAs are usually described as miRNA sponges, although this general scenario may actually rely on few cases [29]. Anyway, this sponge mechanism is not an obvious explanation here because miRNAs are known to foster or repress proliferation and *BRCA* circRNAs should thus only target proliferative miRNAs. Another explanation can be found in the mandatory balance between circRNAs and mRNAs. Assuming that circRNAs biogenesis competes with mRNA, our findings may first sound counterintuitive, as breast tumour tissues would be less buffered by circRNA. It is actually tempting to speculate that this decrease reflects an attempt by the cell to counter the tumour process by lowering the number of circular RNAs in order to maintain a sufficient level of messenger RNAs. Our results echo the recent description of a PTEN circular RNA, circPTEN1, acting as a tumour suppressor downregulated in tumour tissues of colorectal cancer [30] and a previous RNA-seq study showing that circRNAs were globally reduced in tumour tissues from colorectal cancer patients compared to matched normal tissues [31], suggesting a negative correlation of global circRNAs abundance and proliferation. However, this is not supported by our own findings as no significant difference was found between proliferative triple negative subtype tumours and less proliferative luminal subtype tumours. The question remains open whether our findings are restricted to *BRCA1* and *BRCA2* or represents a small part of a global reduction of circRNAs in breast cancer. It cannot be rejected that this reduction in circular RNA abundance possibly reflects a yet unknown tumorigenic hallmark of the tissue, as global splicing dysregulation is known in cancer.

Overall, we developed an innovative assay to study backsplicing and linear splicing, which can be easily translated in diagnostics. We then deciphered the landscape of *BRCA* circRNAs, described novel ones, and demonstrated that the ratio between backsplicing and linear splicing in *BRCA1* and *BRCA2* is not constant but evolve according to the cell environment. At last, we suggested that a disequilibrium between *BRCA1/2* circular and messenger RNAs in favour of mRNA could reflect a tentative adaptation to tumorigenesis. This study should now be pursued on a larger series with other genes from the homologous recombination pathway.

## Supporting information

Supplementary figure 2: Representation of BRCA1 splicing and backsplicing profiles. Exons are drawn as boxes and are numbered, with introns represente

Supplementary figure 3: Representation of BRCA1 splicing and backsplicing profiles. Exons are drawn as boxes and are numbered, with introns represente

Supplementary table 1: BRCA1 and BRCA2 germline splice variations from the validation set. Tissue availability is indicated; in brackets: number of FF

Supplementary table 2: All the BRCA1 circular RNAs in the normal and tumour breast tissue of the validation set. Novel circRNA are indicated in bold.

Supplementary table 3: The novel BRCA1 and BRCA2 circular RNAs in the breast tissue of the validation set.

Supplementary table 4: All the BRCA1 circular RNAs in the normal and tumour breast tissue of the research set. Novel circRNA are indicated in bold.

Supplementary table 5: All the BRCA2 circular RNAs in the normal and tumour breast tissue of the research set. Novel circRNA are indicated in bold.

## Acknowledgements

We thank the Cancéropôle Nord‐Ouest (CNO), Région Normandie, the programme de l’Union Européenne FEDER-FSE/IEJ and the FHU‐G4 genomique for supporting this work. This research study has benefited from the facilities and expertise of the CRB Collection (Breast tissues) of the Henri Becquerel Cancer Center - Rouen - France (https://www.becquerel.fr/le-centre/la-recherche/centre-de-ressources-biologiques).

## Author Contributions

CL conducted most of the biological experiments, participated to design experiments, computational and data analyses and wrote the paper; MV, AD and LB conducted experiments; SC participated to the computational analyses; JCT provided clinical expertise; ML provided pathophysiological expertise; SBD participated to data analyses; PR participated to design experiments, conducted the computational analyses and participated to data analyses; CH conceived, oversaw the research and wrote the paper. All authors have critically read, contributed, edited and approved the final version of this paper.

