## Supplementary figures and images for "Disequilibrium between *BRCA1* and *BRCA2* circular and messenger RNAs plays a role in breast cancer"

### Supplemental Data 1

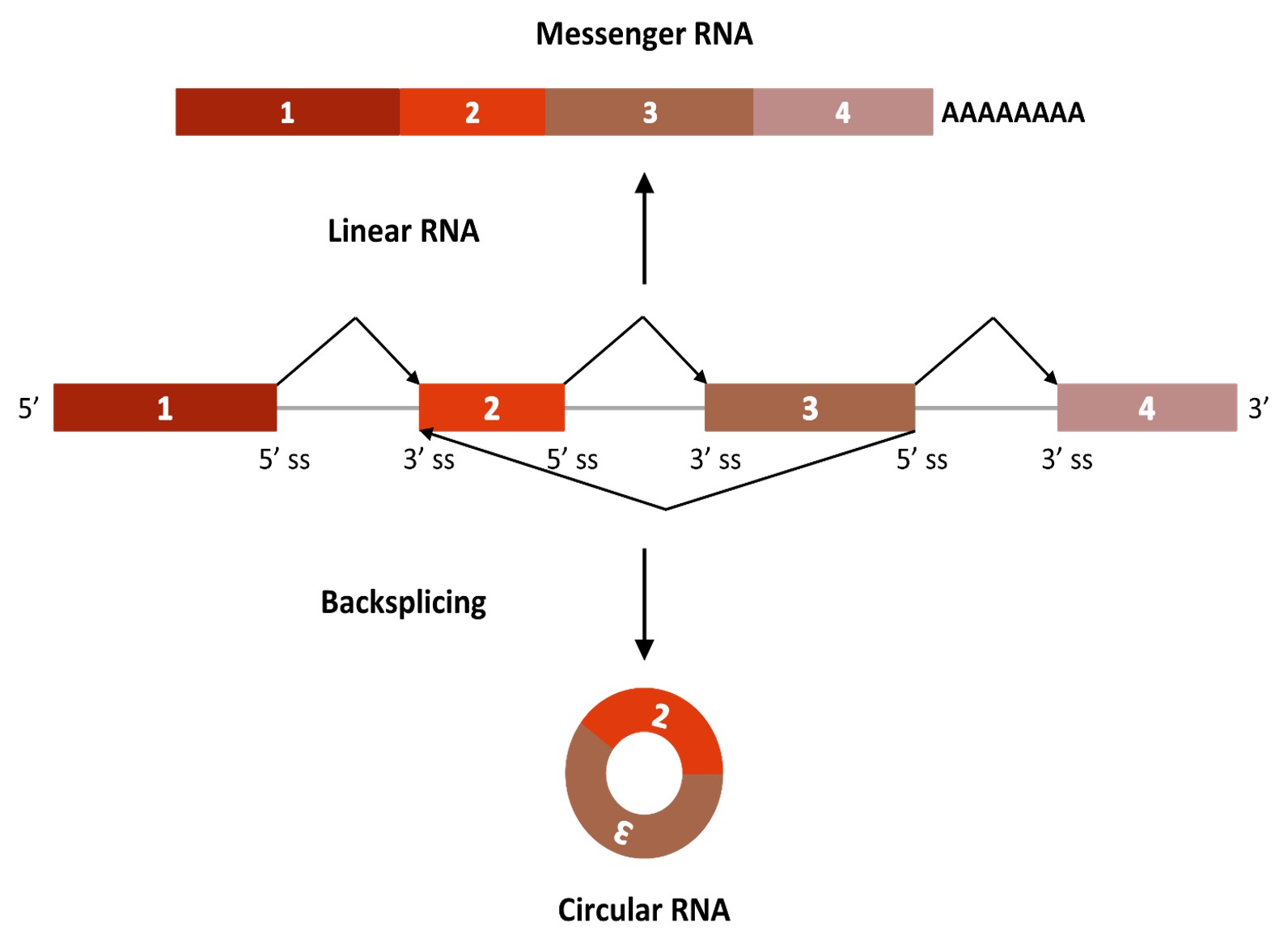

### Supplementary figure 2: Representation of BRCA1 splicing and backsplicing profiles. Exons are drawn as boxes and are numbered, with introns represente

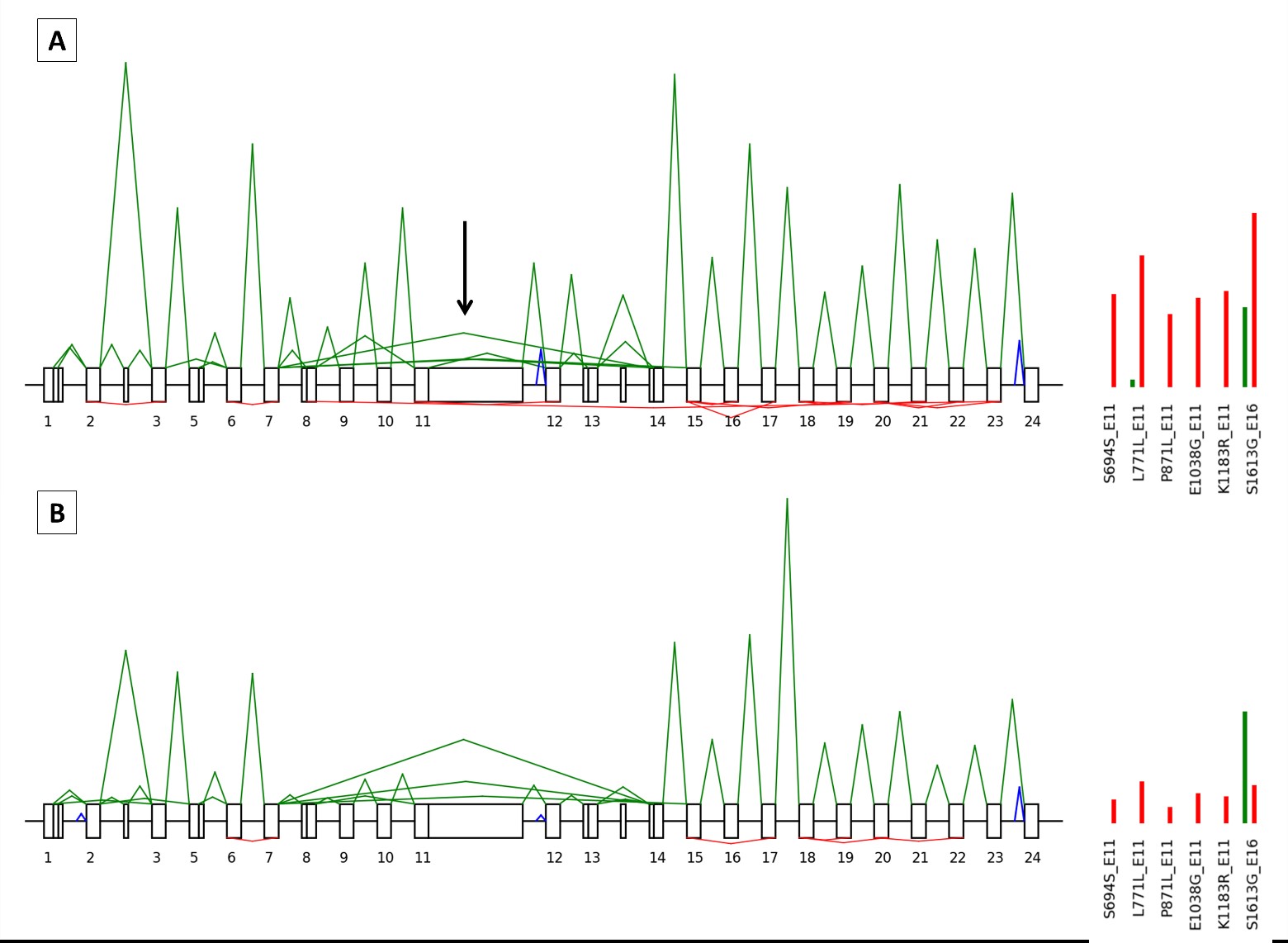

### Supplementary figure 3: Representation of BRCA1 splicing and backsplicing profiles. Exons are drawn as boxes and are numbered, with introns represente

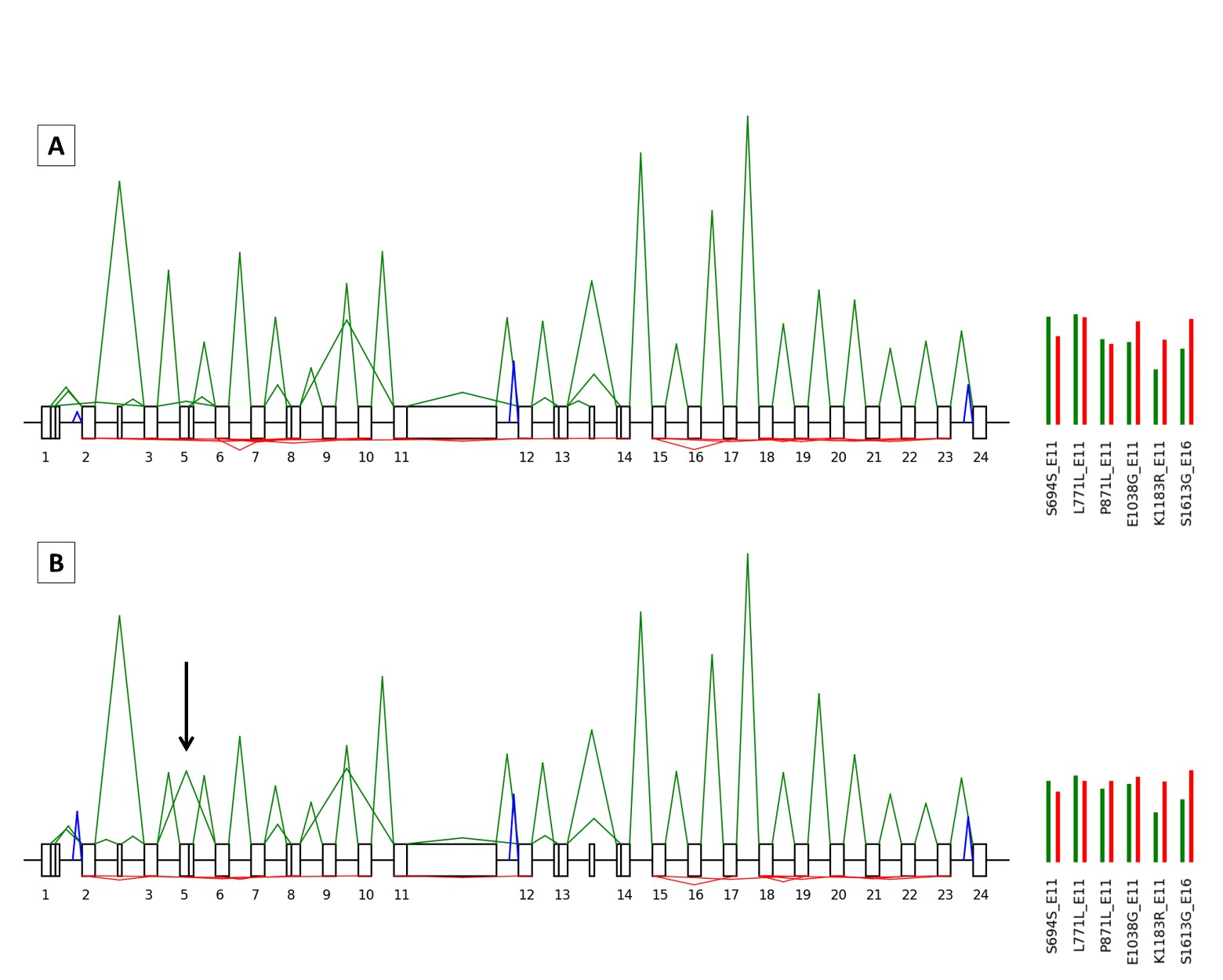
